# Differential LRRK2 signalling and gene expression in WT-LRRK2 and G2019S-LRRK2 mouse microglia treated with zymosan and MLi2

**DOI:** 10.1101/2023.09.14.557532

**Authors:** Iqra Nazish, Adamantios Mamais, Anna Mallach, Conceicao Bettencourt, Alice Kaganovich, Tom Warner, John Hardy, Patrick A. Lewis, Jennifer Pocock, Mark R Cookson, Rina Bandopadhyay

## Abstract

**Introduction:** Mutations in the Leucine Rich Repeat Kinase 2 (*LRRK2*) gene cause autosomal dominant Parkinson’s disease (PD) with the most common causative mutation being the *LRRK2* p.G2019S within the kinase domain. LRRK2 protein is highly expressed in the human brain and also in the periphery, and high expression of dominant PD genes in immune cells suggest involvement of microglia and macrophages in inflammation related to PD. LRRK2 is known to respond to extracellular signalling including TLR4 resulting in alterations in gene expression, with the response to TLR2 signalling through zymosan being less known.

**Methods:** Here, we investigated the effects of zymosan, a TLR2 agonist and the potent and specific LRRK2 kinase inhibitor MLi-2 on gene expression in microglia from *LRRK2-WT* and *LRRK2* p.G2019S knock-in mice by RNA-Sequencing analysis.

**Results:** We observed both overlapping and distinct zymosan and MLi-2 mediated gene expression profiles in microglia. At least two candidate Genome-Wide Association (GWAS) hits for PD, CathepsinB (*Ctsb*) and Glycoprotein-nmb (*Gpnmb*), were notably downregulated by zymosan treatment. Genes involved in inflammatory response and nervous system development were up and downregulated respectively with zymosan treatment while MLi-2 treatment particularly exhibited upregulated genes for ion transmembrane transport regulation. Furthermore, we observed the top twenty most significantly differentially expressed genes in *LRRK2* p.G2019S microglia show enriched biological processes in iron transport and response to oxidative stress.

**Discussion:** Overall, these results suggest that microglial LRRK2 may contribute to PD pathogenesis through altered inflammatory pathways. Our findings should encourage future investigations of these putative avenues in the context of PD pathogenesis.

## 1. Introduction

The etiopathology of PD is multifactorial with a complex interplay of genes and environmental triggers^1,2^. Over the past decades, research has increasingly provided evidence that links neuroinflammation to neurodegeneration in PD. For example, microglia positive for human leukocyte antigen D (HLA-DR) have been identified in the substantia nigra pars compacta of PD patients^3^. Furthermore, there is an increase in inflammatory biomarkers including tumour necrosis factor alpha (TNFα), interleukin (IL)-1β, IL-6, epidermal growth factor (EGF) and transforming growth factor alpha (TGFα) in the brain and cerebrospinal fluid (CSF) of PD patients^4,5^. Similarly, increased IL-1β, IL-6 and gamma delta+ T cells, implicated in inflammation, occurs in blood and brain of PD patients^6,7^, supporting the potential involvement of immunological events in neurodegeneration.

Mutations in the *LRRK2* gene cause autosomal dominant disease with clinical features that are indistinguishable from sporadic PD. The majority of the well characterized pathogenic mutations in *LRRK2* are found in the two active catalytic domains (GTPase and kinase) of the encoded large multidomain protein. The common *LRRK2* p.G2019S mutation is located within the kinase domain and increases LRRK2 kinase activity^8^. LRRK2 protein is expressed within the brain and in peripheral tissues with notably high expression in microglia and macrophages^9^. High expression of dominant PD genes within microglia and/or macrophages suggests that inflammation is relevant for PD development. Knockdown, knockout or pharmacological inhibition of LRRK2 functions results in a decrease in inflammatory responses^10,11^, further highlighting a potential role of LRRK2 in immune function^12^. Based on these observations, LRRK2 has been identified as potential therapeutic target for anti-inflammation strategies in PD^13^

Inflammatory pathways are typically organized where cell surface receptors respond to external signals to trigger intracellular signalling events that result in alterations in gene expression. Prior work has shown that LRRK2 is responsive to extracellular signalling including TLR4^14^. Here, we have investigated the effects of the TLR2 agonist zymosan on gene expression in microglia from *LRRK2* p.G2019S knock-in mice by RNA-Seq analysis. We demonstrate that zymosan triggered significant upregulation in LRRK2 phosphorylation which remained significantly inhibited with MLi-2 treatment. We also show that the two genotypes, wild-type and G2019S-*LRRK2*, and two treatments, zymosan and MLi-2, induced overlapping as well as distinct effects on gene expression profile of microglial cells. Zymosan consistently displayed a stronger effect on gene expression as compared to MLi-2 treatment and G2019S-LRRK2 genotype.

## 2. Materials and Methods

### 2.2. Animals and primary microglia cultures and treatment

Homozygous *LRRK2* G2019S knock-in mice^15,16^ and C67Bl/6J wild-type mice were housed at NIH, and all animal procedures were carried out in strict accordance with the guidelines provided by the Care and Use of Laboratory Animals of NIH as approved by the Institutional Animal Care and Use Committees of the US National Institute on Aging (Approval number: 463-LNG-2018).

Primary mixed glial cultures were established from the brains of wild-type or from knock-in mice harbouring G2019S-*LRRK2* mutation at postnatal days 1-2 (P1-2). After approximately 10 days, primary microglia were shaken from mixed glial cultures as described in Russo et al^15^. Purity of the obtained culture was verified using double immunofluorescence method by staining the culture with rabbit anti-Iba1 (Abcam #ab5076) for microglia, rabbit anti-Glial Fibrillary Acidic Protein (GFAP) for astrocytes (Dako Omnis #GA524) and anti-4′,6-diamidino-2-phenylindole (DAPI) (Cell Signaling #4083). The primary microglial yield was approximately 5 × 10^5^ cells/flask with minimal astrocyte contamination. For immunoblotting, microglia were seeded and treated with pro-inflammatory agents for 4 hours and 24 hours. For RNA-Seq, cells were seeded and treated with pro-inflammatory agents for 24 hours only. For immunoblottting, cells were lysed with the lysis buffer consisting of 10% cell lysis buffer (10x) (Cell Signaling #9803S), 1% protease inhibitor cocktail (100x) (Cell Signaling #5871S) and 1% phosphatase inhibitor cocktail (100x) (Cell Signaling #5870S), centrifuged at 3200g for 5 min, collected and stored at -80°C for future use. For RNA-Seq collection, RNA was extracted from the primary microglia using TRIzol reagent (Invitrogen #15596026) as per manufacturer’s protocol.

### 2.2. Protein assay

Protein levels were measured using the Thermofisher Pierce BSA Standard Pre-Diluted Set as per manufacturer’s instructions using BSA as standard.

### 2.3. Zymosan and MLi-2 treatment protocol

Zymosan A from *Saccharomyces cerevisiae* (Sigma #Z4250) and MLi-2 from Merck (Tocris #5756) were used for these experiments as TLR2 inflammatory stimuli and LRRK2 kinase inhibitor respectively. For immunoblots, primary microglia from wild-type and G2019S-LRRK2 Tg mice were treated with 200μg/ml zymosan and 1μM MLi-2 for 4 hours and 24 hours, and for RNA-Seq, they were treated for 24 hours only.

### 2.4. Immunoblots and fluorescent blots

Samples were heated to 95°C for 3 minutes in loading buffer NuPAGE LDS Sample Buffer (4x) (Invitrogen NP0008) with 5% β-Mercaptoethanol then loaded on 4-20% Criterion TGX Precast Midi protein gels (Bio-Rad #5671094). The gels were run in TGS buffer (1x) and transferred using Trans-Blot Turbo Mini 0.2μm Nitrocellulose Transfer Packs (Bio-Rad #1704158) using Trans-Blot Turbo Transfer System (Bio-Rad). Membranes were incubated with primary antibodies overnight and β-actin for 1 hour. After the membranes were incubated with the appropriate Fluorescent-conjugated secondary antibodies, protein bands were visualised using Odyssey CLx (Li-Cor) and quantified using Bio-Rad Lab Image (version 6.1) software.

### 2.5 RNA Sequencing (RNA-Seq)

RNA quality and integrity was measured using the Agilent 2100 Bioanalyzer RNA 6000 Nano Chip (Agilent) and analysed on 2100 Expert Software. The samples were then stored at -80°C until further use. The RNA-Seq was carried out by Psomagen labs, whole-genome sequencing service providers in USA, through Illumina SBS technology using the Total RNA Ribo-Zero Gold Library Kit.

### 2.6. Analysing RNA-Seq data

RNA-Seq reads were aligned to the mouse reference genome (mm10) using STAR^17^ and expression counts per transcript was quantified using eXpress^18^ followed by using DESEQ2^19^ to normalise data. Gene expression data was loaded into MATLAB version R2019b to identify up- and downregulated genes in different treatment and genotype groups through differential gene expression analysis. Subsequently, functional enrichment analysis tools FunRich version 3.1.3 and Hippie version 2.2 were used to perform functional enrichment and interaction network analysis on the identified up- and downregulated genes in different treatment groups, mapping genes

### 2.7. Statistical analyses

One-way analysis of variance (ANOVA) with Tukey’s post-hoc test was used to compare the difference between the means of three or more treated and untreated independent groups. Unpaired two-tailed student’s t-tests were used to compare the difference between the means of any two normally distributed groups e.g. when determining the difference between non-stimulated cells and cells stimulated with zymosan. The statistical tests used for each data set are mentioned where appropriate in the legends of every experimental figure. All data were analysed using GraphPad Prism 5, R Studio version 1.3.1093, MATLAB version R2019b. All quantitative data are expressed as mean ± S.E.M. and represent in most cases at least three independent set of experiments. The exact number of experiments with internal replicates is mentioned in the legends of every experimental figure. Statistical significance was set at p < 0.05.

## 3. Results

### 3.1. Zymosan induces phosphorylation of LRRK2 in microglia

LRRK2 is phosphorylated at a series of residues between the ankyrin and leucine-rich repeat domains, including Serine935^20^. Phosphorylation of LRRK2 at S935 is controlled by other kinases, notably casein kinase 1α^21^, but is also responsive to inhibition of LRRK2 kinase activity^22^. We therefore used pS935 as a proxy for LRRK2 activation. A significant upregulation of LRRK2 phosphorylation at Ser935 was observed with zymosan treatment at 4 hours and 24 hours (Figure 1) in wild-type or G2019S knock-in cells. Co-treatment of cells with zymosan and MLi-2 resulted in a significant decrease in pS935-LRRK2 levels at 4 hours in both genotypes and at 24 hours in wild type cells.

**Figure 1.**
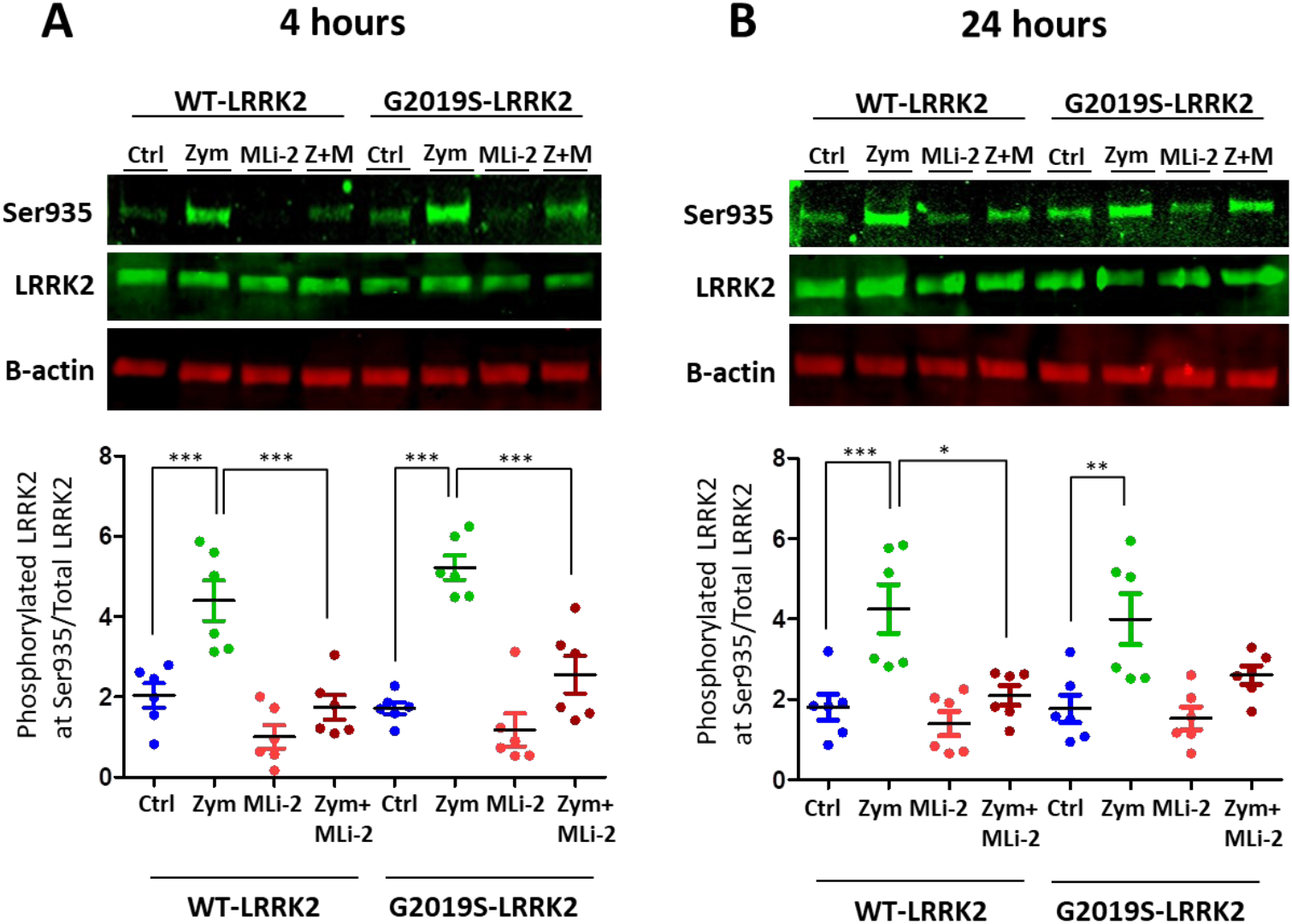
LRRK2 phosphorylation evoked by zymosan is inhibited with MLi-2 treatment in wild-type and G2019S-LRRK2 microglia. Wild-type and G2019S-LRRK2 were treated with 1μM MLi-2 and 200μg/ml zymosan for 4h and 24h. Fluorescent immunoblots and corresponding quantifications are shown for (A) 4h and (B) 24h. Blots were probed with LRRK2 phosphorylation antibody pSer935 and total LRRK2. Controls contain media only. Values represent the mean ± S.E.M. of 2 independent experiments (with internal triplicates in each experiment). * and ** and *** signify p<0.05, 0.01 and 0.001, respectively. Colours for graphs indicate: Blue: Control, Green: zymosan, Red: MLi-2, Dark red: zymosan+MLi2 treatments. Ctrl: Control, Zym/Z: zymosan, M: MLi-2

### 3.2. Differential gene expression between wild-type and G2019S-LRRK2 with zymosan and MLi-2

Having established that zymosan treatment results in an increase in pS935 LRRK2, we next performed RNA-Seq to evaluate the effects of LRRK2 modulation on inflammatory signalling in two genotype groups (WT and G2019S) and four treatments (control, MLi-2, zymosan and zymosan plus MLi-2). Mean vs variance plots for gene expression after variance stabilizing transformation indicate that variance was consistent across gene expression levels in both genotypes (Supplementary Figure S1A,B).

We next examined the first two principal components of the overall gene expression patterns against genotype and treatment groups. PC1, which explains ∼90% of the variance, aligns with zymosan treatment, with or without the addition of MLi-2, while PC2, which captures ∼ 5% of the variance, appears to be largely explained by genotype (Figure 2). Consistent with this overall view of the data, clustering of overall gene expression by Euclidean distance show a clear separation between zymosan treated samples and samples without zymosan, followed by separation by genotype and finally MLi-2 treatment (Figure 3A). Two additional heatmaps (Figure 3B) reveal the top twenty differentially expressed genes with the highest mean expression across all genotypes and treatments.

**Figure 2.**
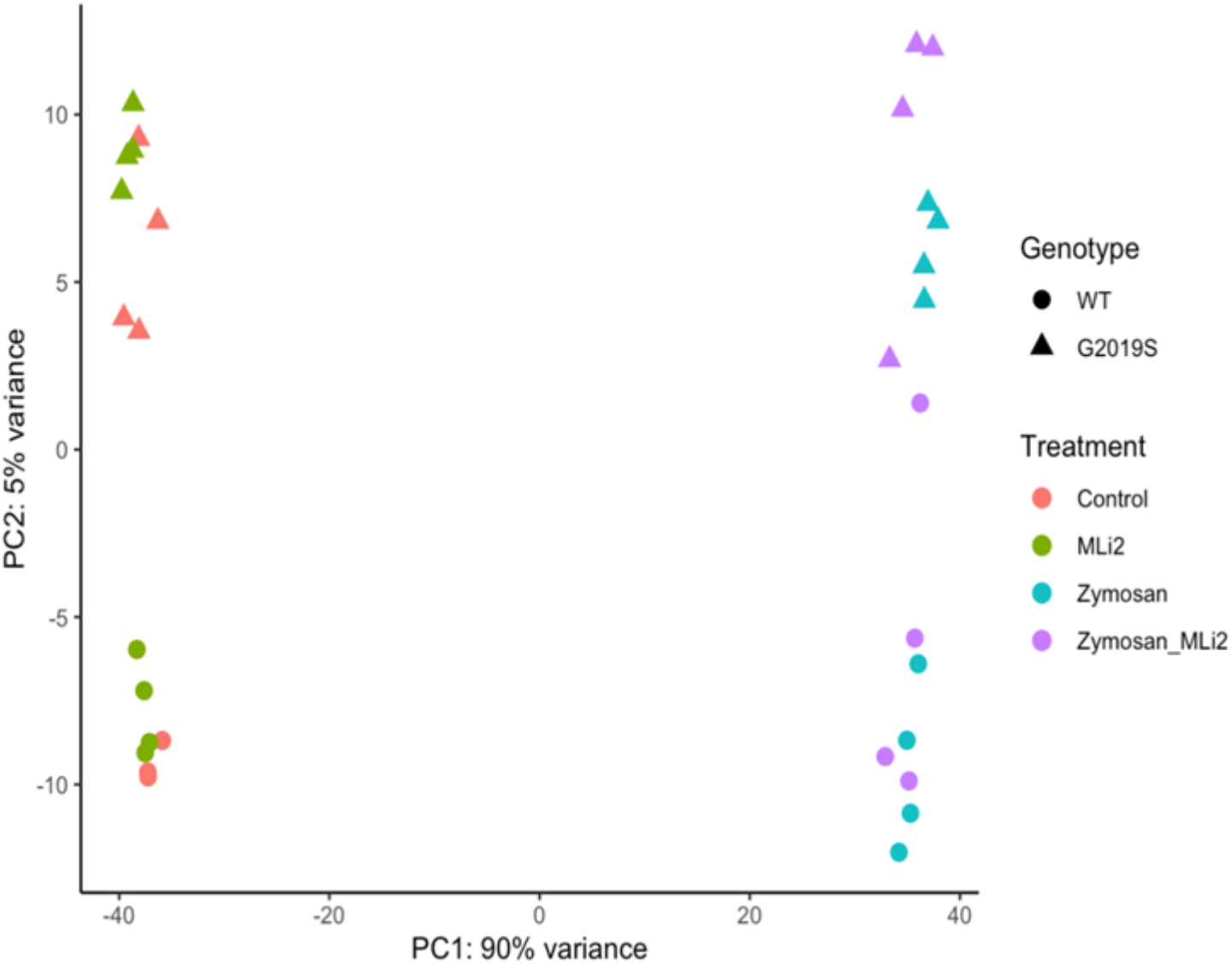
RNA-Seq profiling showed by first two principal components of wild-type and G2019S-LRRK2 microglia after zymosan and MLi-2 stimulation. First two principal components, PC1 and PC2, of all samples used in the study. Data contains two biological variables, treatment with zymosan and MLi-2 and genotype (wild-type and G2019S-LRRK2). Note that samples separate largely by treatment and to a lesser extent by genotype.

**Figure 3.**
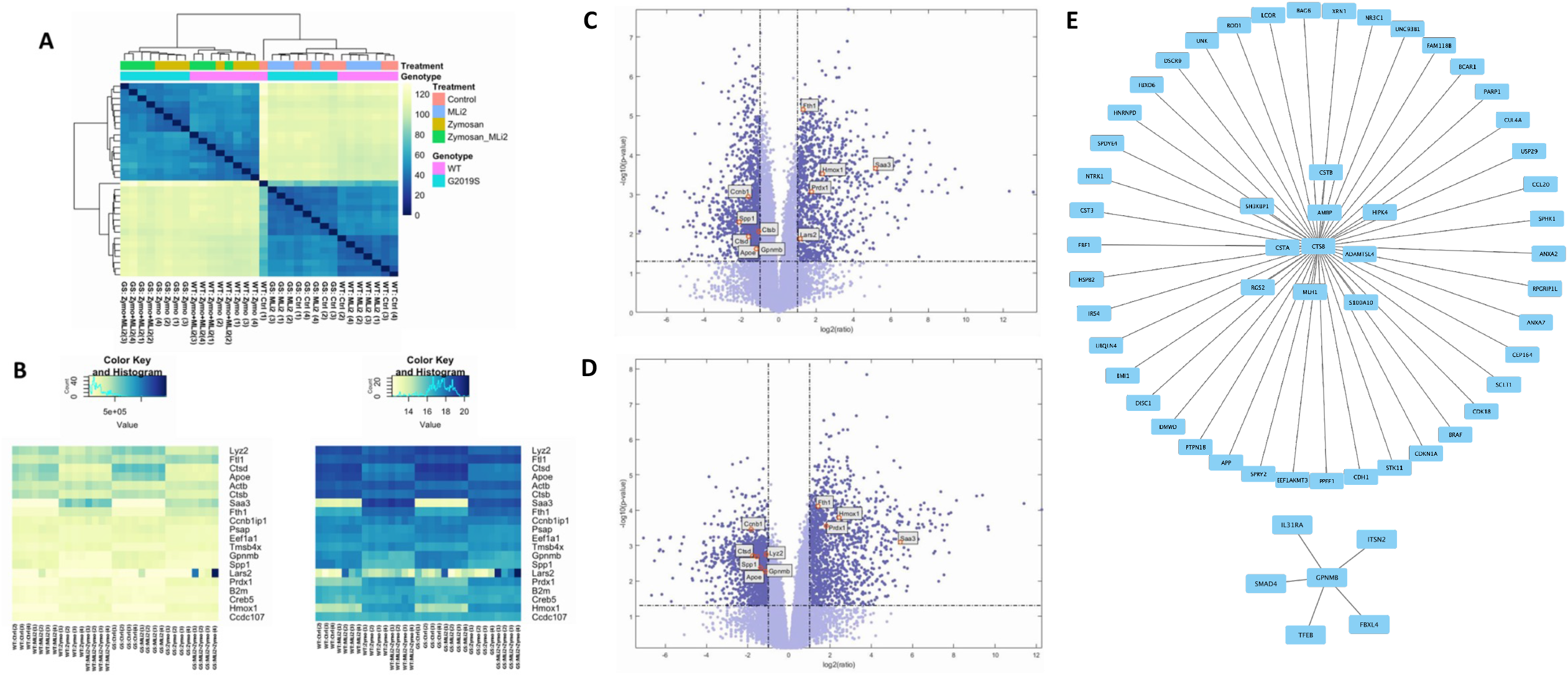
Heatmaps and clustering analysis of RNA-Seq profiling of wild-type and G2019S-LRRK2 microglia after zymosan and MLi-2 stimulation, Volcano plots highlighting significantly differentially expressed genes with zymosan treatment in wild-type and G2019S-LRRK2 genotypes and Protein interaction maps. (A) Hierarchical clustering and heatmap of all samples used in this study. Colours in the heatmap represent the Euclidean distance between samples in a pairwise manner, scaled as shown on the upper right yellow-blue scale. Above the heatmap is a colour representation of the model variables, which included two biological variables, treatments with zymosan and MLi-2 and genotype (wild-type and G2019S-LRRK2). Note that samples separate largely by treatment and to a lesser extent by genotype. (B) Heatmaps for the top twenty most statistically significant genes with the highest mean expression across all samples. Each gene on the right side of each heatmap is coloured by Z (normalized standard deviations from the mean) for expression relative to the overall mean expression for that gene and samples are listed below each heatmap. (C) shows significantly differentiated genes with zymosan treatment in G2019S-LRRK2 and (D) shows genes in wild-type. Each point represents a significantly differentiated gene. Dark blue colour depicts genes which passed the thresholds for 2-Log Fold Change with upper right hand quadrant showing upregulated and upper left hand quadrant showing downregulated genes. The top twenty genes with the highest mean expression across all samples shown in (B) are shown in boxes in these volcano plots. (E) Protein interaction maps generated with Hippie software showing the interaction of Ctsb protein (blue) and Gpnmb protein (green) with other interacting proteins. Ctrl: Control, Zymo: Zymosan

We next examined gene expression between treatment groups. Volcano plots show significantly differentially expressed genes with zymosan treatments in both wild-type and G2019S-LRRK2 genotypes (Figures 3C and 3D). There were abundant differences driven by zymosan treatment including *Fth1* and lysosomal genes, which we have previously shown to be responsive to LPS-induced inflammation in microglia^23^. We also note that at least two candidate GWAS hits for PD, namely *Ctsb*) and *Gpnmb*, are downregulated by Zymosan treatment, irrespective of genotypes.

### 3.3. Functional enrichment analysis in wild-type and G2019S-LRRK2 with zymosan and MLi-2

Two protein interaction maps generated with Hippie software show the interaction of *Ctsb* and *Gpnmb* proteins with other interacting proteins. Ctsb and Gpnmb from the list of top twenty genes (Figure 3E) are also GWAS hits for PD^24^ hence it is interesting to see their interacting proteins in the context of PD.

Ontological analysis of significant differentially expressed genes was performed using FunRich software (version 3.1.3). Figure 4 shows biological processes (A-B) and cellular components (C-D) significantly enriched in zymosan-treated wild-type and G2019S-LRRK2 microglia. The most enriched GO terms for biological processes in upregulated genes were inflammatory response and nervous system development in downregulated genes. The most enriched GO terms for cellular components were plasma membrane for upregulated genes and glutamatergic synapse for downregulated genes.

**Figure 4.**
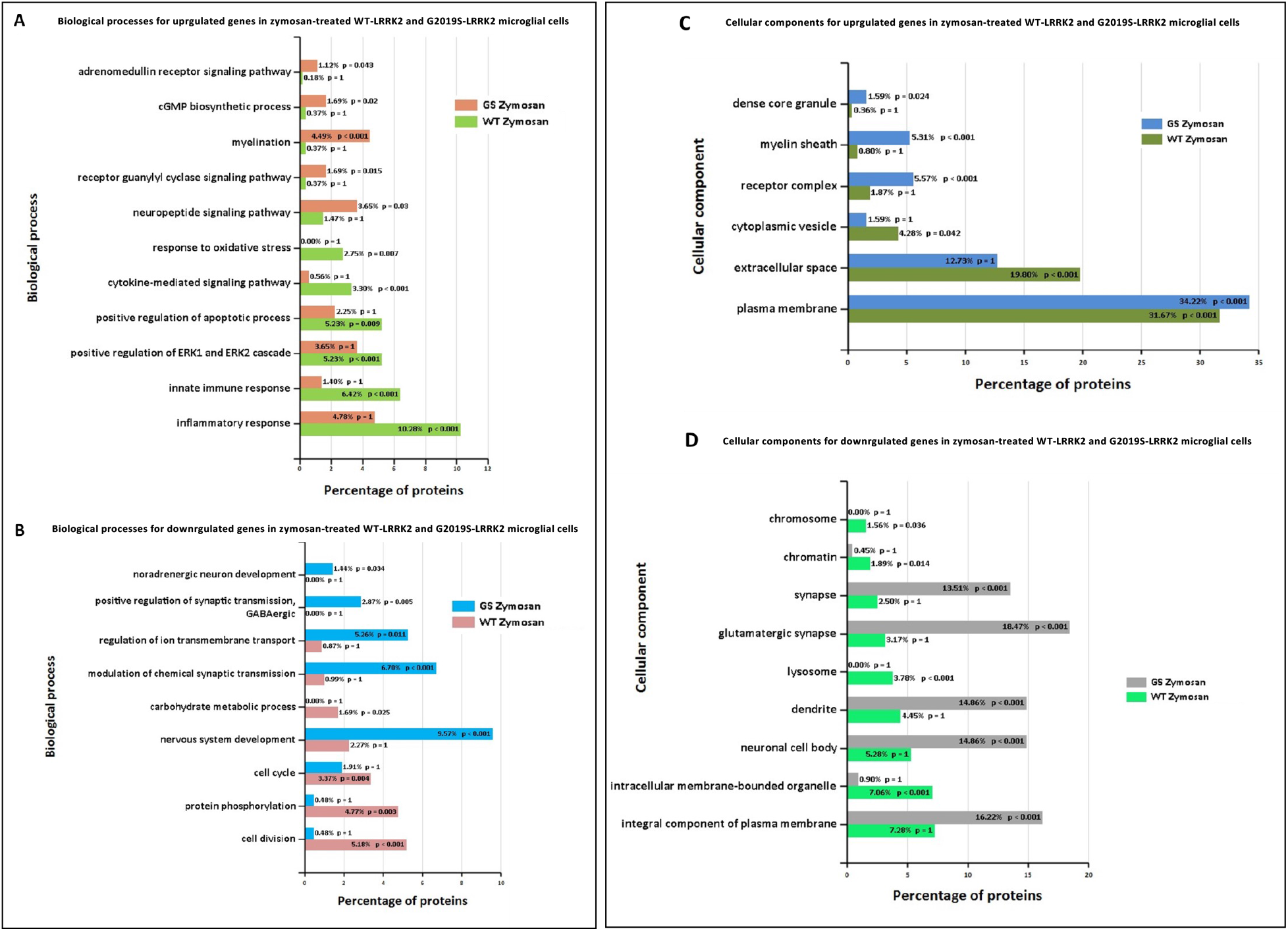
GO enrichment analysis of differentially expressed genes in zymosan-treated wild-type and G2019S-LRRK2 microglia. GO (A-B) biological process and (C-D) cellular component enrichment analyses of (A,C) upregulated proteins and (B,D) downregulated proteins in zymosan-treated wild-type and G2019S-LRRK2 microglial cells were performed using FunRich functional enrichment analysis tool. Significantly enriched GO terms are shown with Benjamini-Hochberg and Bonferroni-corrected p-values. Statistical significance was taken at p < 0.05

Furthermore, Figure 5 show (A-B) biological processes and (C) cellular components significantly enriched in MLi-2 treated wild-type and G2019S-LRRK2 microglia. The most enriched GO terms for biological processes in upregulated genes were regulation of ion transmembrane transport and positive regulation of apoptotic process in downregulated genes. The most enriched GO terms for cellular components were plasma membrane for downregulated genes and no significantly enriched cellular components terms for upregulated genes (Supplementary Table S1). Notably, Figure 6 shows (A) biological processes and (C) cellular components significantly enriched for top twenty most significantly differentially expressed genes in G2019S-LRRK2 microglia. The most enriched GO terms for biological processes in upregulated genes were iron ion transport and response to oxidative stress and extracellular space for the most enriched GO terms for cellular components.

**Figure 5.**
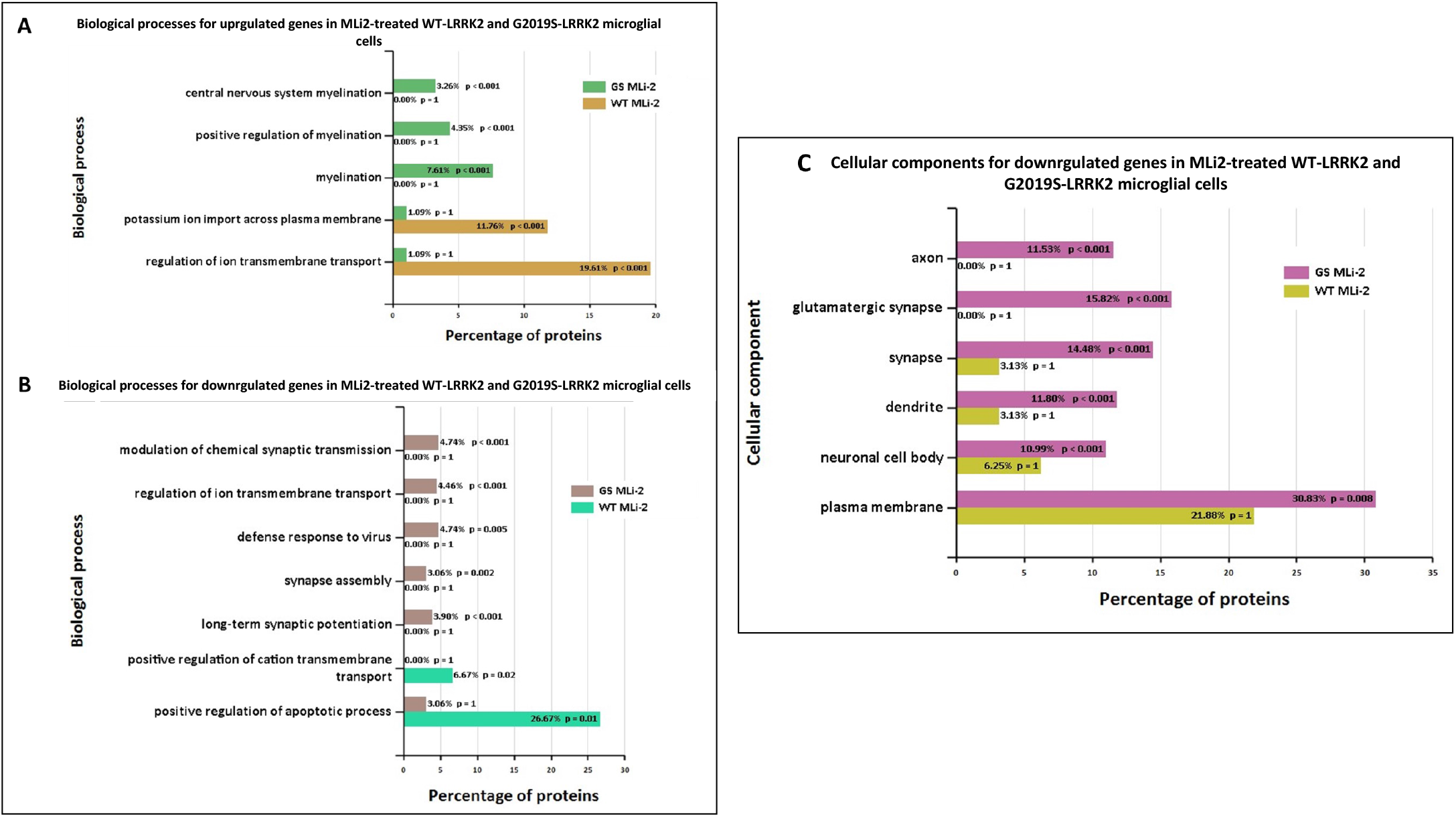
GO enrichment analysis of differentially expressed genes in MLi2-treated wild-type and G2019S-LRRK2 microglia. GO (A-B) biological process and (C) cellular component enrichment analyses of (A) upregulated proteins and (B,C) downregulated proteins in MLi2-treated wild-type and G2019S-LRRK2 microglial cells were performed using FunRich functional enrichment analysis tool. There were no significantly enriched GO terms with upregulated proteins in MLi2-treated wild-type and G2019S-LRRK2 microglial cells in my data, hence no data for that is shown. Significantly enriched GO terms are shown with Benjamini-Hochberg and Bonferroni-corrected p-values. Statistical significance was taken at p < 0.05

**Figure 6.**
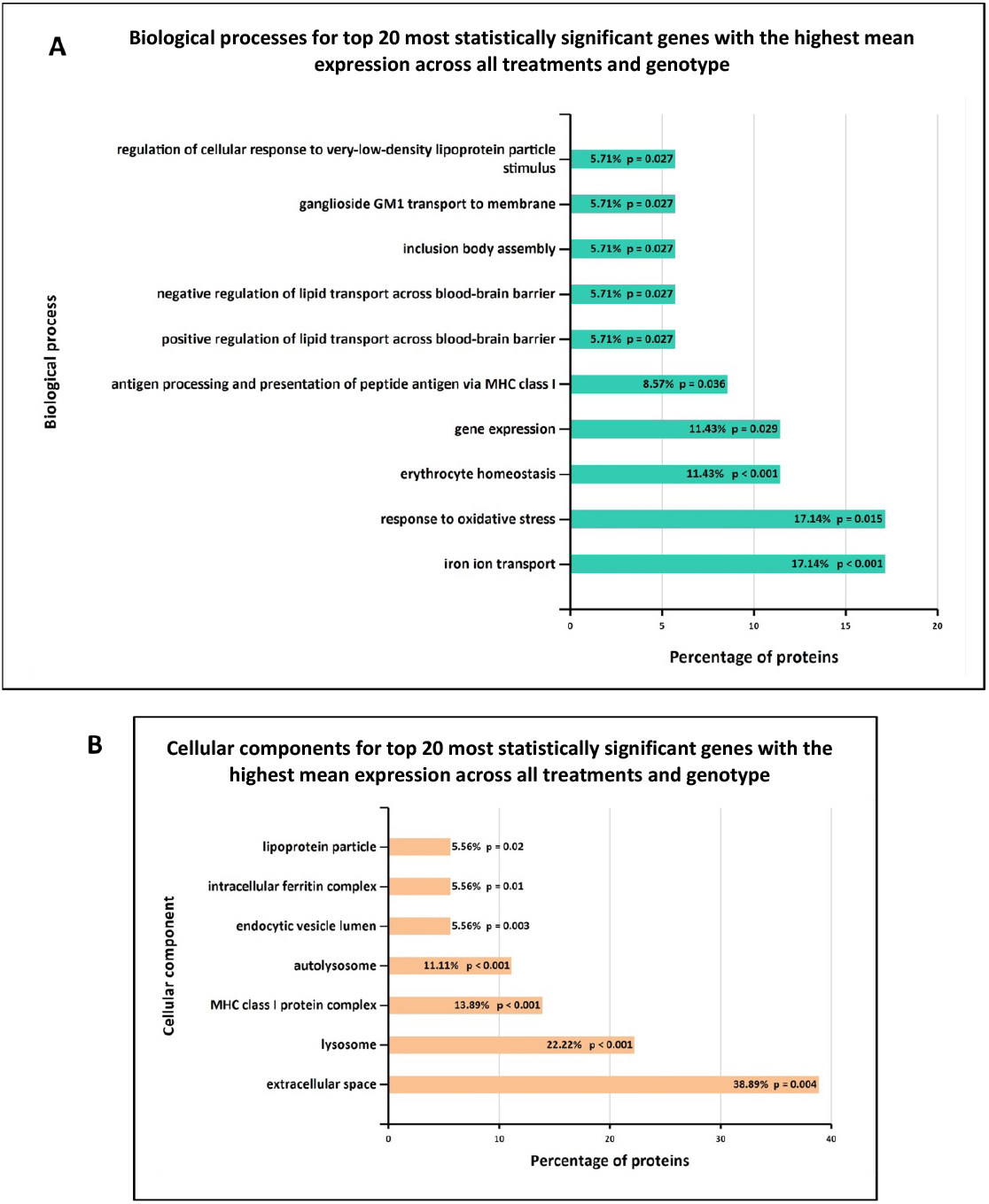
GO enrichment analysis of the top twenty most statistically significant genes with the highest mean expression across all treatments and genotype. GO (A) biological process and (B) cellular component enrichment analyses of top twenty most statistically significant differentially expressed genes across all samples (from Figure 5) were performed using FunRich functional enrichment analysis toolSignificantly enriched GO terms are shown with Benjamini-Hochberg and Bonferroni-corrected p-values. Statistical significance was taken at p < 0.05

## 4. Discussion

Microglia are resident innate immune cells of the CNS, contributing neuroprotective characteristics during acute immune responses but are also implicated as mediators of cell loss in neurodegenerative disorders after chronic activation. Activated microglia have been found in SNpc of PD patients^25^ and are also implicated in causing damage to dopaminergic neurons^26^. The active role of microglia in neuroinflammation is also reviewed in a study by Perry et al^27^. Upon activation by inflammatory stimuli, microglia switch from resting to activated state, releasing pro-inflammatory and reactive oxygen species to mediate an inflammatory response^28^. LRRK2 is highly expressed in microglia^9^, indicating a possible role for LRRK2 and microglia in contributing to PD pathogenesis through altered inflammatory signalling. The inflammatory stimulus LPS is widely used to trigger microglial activation as previously carried out^14^, and are used to analyse the transcriptomic profile of primary microglial cells by various studies^29,30^. To our knowledge, transcriptomic data using zymosan as an inflammatory stimulus in the context of LRRK2 has not been investigated to date.

To dissect novel LRRK2-related biological processes in microglia, we performed RNA-Seq analysis of wild-type and G2019S-LRRK2 cells under basal conditions, or after stimulation with zymosan and in the presence of the LRRK2 kinase inhibitor, MLi-2. We observed various overlapping and individual effects on gene expression and induction of biological processes. Our data has shown zymosan to have a stronger effect on gene expression as compared to MLi-2 treatment or genotype. This was confirmed with first principle component of the overall gene expression profile which separated zymosan from MLi-2 and basal treatment groups, whereas genotype had a more subtle effect on overall gene expression. The results are unsurprising given that zymosan is a strong stimulator of immune signalling similar to LPS with LRRK2 likely playing a modulatory role through altered lysosomal responses^23^.

Enrichment analyses showed that some of the most enriched GO terms for biological processes in zymosan-treated wild-type microglial cells were inflammatory response, innate immune response and cytokine-mediated signalling pathways (Figure 4A). However, these GO terms were not significantly enriched in G2019S genotype, which is suggestive of the potential involvement of G2019S-LRRK2 (hyperkinetic) in inflammatory responses. Pathways significantly enriched with zymosan treatment in G2019S genotype involved adrenomedullin receptor signalling pathway, cyclic guanosine monophosphate (cGMP) biosynthetic process, receptor guanylyl cyclase signalling pathway and neuropeptide signalling pathway which were not observed in wild-type cells (Figure 4A). Therefore, it may be important to investigate these pathways in the context of PD pathogenesis. Adrenomedullin has been recently discussed to be an important participant in neurological diseases and is seen to exhibit neuroprotective effect against brain insults^31^. Similarly, guanylyl cyclase-cGMP signalling and the neuropeptides and neurotransmitters are also known to play a role in PD pathogenesis^32,33^. GO enriched terms also indicate zymosan to be a positive regulator of ERK1 and ERK2 signalling cascade and apoptotic processes. ERK kinases belong to the MAPK superfamily of kinases which are known to be implicated in PD^34^. The most significantly enriched terms for biological processes for downregulated proteins with zymosan treatment in G2019S genotype cells include nervous system development, regulation of ion transmembrane transport and modulation of chemical synaptic transmission indicating these processes to perhaps be negatively affected in PD.

In terms of cellular processes, the most enriched terms for zymosan treatment in wild-type and G2019S genotype include plasma membrane, extracellular space and glutamatergic synapse. MLi-2 on the other hand seemed to have a more subtle effect on expressed genes and enriched GO terms. Interestingly, for enriched biological processes in G2019S, upregulated genes in MLi-2 seemed to only significantly enrich processes involved in myelination. However, the underlying mechanisms of the inflammatory effects and role of microglia is yet to be fully understood in the context of PD.

Furthermore, in basal G2019S condition, downregulated proteins were significantly enriched for chemotactic pathways such as natural killer cell, lymphocyte and eosinophil chemotaxis. This is suggestive of suppressed chemotactic abilities in PD pathogenesis and indeed a recent study has found lymphocyte chemotaxis to be impaired in PD patients^35^. Furthermore, it also suggests that microglia expressing G2019S have an inability to become recruited to areas of damage within the PD brain parenchyma. A similar microglial response is linked to the AD risk gene TREM2^36^. Defence response to virus was also significantly enriched for downregulated proteins in G2019S genotype which is confirmative of impaired response to virus as one of the key contributors in PD pathogenesis. Previous studies have shown that systemic and local inflammatory responses to viruses have potential involvement in neuronal damage, even in the absence of neuronal cell death. Viruses are seen to evoke CNS inflammation either by entering the brain or by crossing a damaged BBB or along the peripheral nerves or by activating the peripheral innate and adaptive immune responses^37,38^. Cellular components most significantly enriched in upregulated proteins in G2019S genotype also consist of plasma membrane, myelin sheath and internode and paranode regions of axon, rendering investigating all these biological processes and cellular components important in the context of PD^39,40^.

The top twenty genes which showed the highest mean expression across all genotype and treatment conditions also revealed some intriguing aspects to the results. First, two of the twenty genes, *Gpnmb* and *Ctsb*, were reported as genetic risk loci for PD in a recent GWAS study^24^. Elevated expression of *Gpnmb* is associated with increased PD risk^41^ and through its interaction with alpha-synuclein, it may also confer increased disease risk^42^. It is noteworthy that CathepsinD (*ctsd*) gene is mutated in human neuronal ceroid lipofuscinosis ^43,44^. Interestingly, LRRK2 *G2019S* leads to suppression of lysosomal proteolytic activity in macropahges and it regulates the abundance of multiple lysosomal proteins^45^. Therefore, research on Ctsb, Ctsd and Gpnmb in relation to LRRK2 and their interacting proteins should be expanded in order to further elucidate the molecular underpinnings of PD. Enriched biological processes for the top twenty genes show most significantly enriched terms to be iron ion transport and response to oxidative stress and enriched cellular components show extracellular space and lysosomes to be most significantly enriched (Figure 6). These are indeed important biological processes and cellular components implicated in PD.

## Summary/Conclusion

With this data, our study is the first to report the identification of differential gene expression and dissection of novel biological pathways in response to zymosan and LRRK2 kinase inhibition with MLi-2 in G2019S-LRRK2 KI primary microglia cells. The data presented in this paper raise several interesting points and opens up new pathways of PD research that would be suitable for further investigation. In this study, we assessed LRRK2 activation *via* phosphorylation of Ser935 – an indirect measure, mediated by an upstream kinase. In future work, it will be important to assess direct measures of LRRK2 kinase activity such as LRRK2 Ser1292 phosphorylation ^46^, and phosphorylation of Rab proteins such as Rab10, Rab12 or Rab8a. One interesting future research direction could be to utilize human-derived cells with native LRRK2 expression. It would be necessary to understand whether human primary cells have the same phenotype as the mouse equivalents, or whether they diverge in phenotype. With the increasing use of stem cell derived human cells to model disease, assessing zymosan induced changes in expression in pluripotent stem cell derived macrophages and microglia with and without G2019S-LRRK2 mutations will be insightful for a deeper, and more human relevant, analysis. As noted above, the various enriched biological pathways discussed in this paper should be highlighted and prioritised for investigation in the context of PD pathogenesis – especially the regulation of genes implicated by genome wide association studies for PD that are linked to lysosomal function. Finally, it is hoped that the findings from the research conducted for this paper will help advance the understanding of PD pathogenesis and drug development towards new therapies for the millions of people world-wide living with PD^47^.

## Legends

Supplementary Figure 1: Standard deviation to visualise the variance of the data. The plots show the standard deviations of the transformed data, across all samples, against the mean, using the shifted logarithm transformation, the regularized log transformation and the variance stabilizing transformation. Each point represents a differentially expressed gene and (A) shows variance between this differential expression between genes in G2019S-LRRK2 and (B) shows variance in wild-type.

Supplementary Table S1. List of RNA Seq transcripts up and down regulated in WT and G2019S mice with Zymosan and MLi2 treatment.

## Acknowledgements

RB is funded by the Reta Lila Weston Trust for Neurological Studies. IN was a recipient of the Yule Bogue Fellowship (UCL) to carry out the experiments at NIA/NIH. This work was supported in part by the Michael J. Fox Foundation and the National Institutes of Health grants NS110188 and AG077269 to Adamantios Mamais. Anna Mallach was supported by BBSRC London Interdisciplinary Biosciences Consortium (LIDo; BB/M009513/1). CB is supported by Alzheimer’s Research UK (ARUK-RF2019B-005) and Multiple System Atrophy Trust.

## Notes

### Competing Interest Statement

The authors have declared no competing interest.

