## Supplementary figures and images for "Differential LRRK2 signalling and gene expression in WT-LRRK2 and G2019S-LRRK2 mouse microglia treated with zymosan and MLi2"

### IN Suppl Fig S1

#### Slide 1
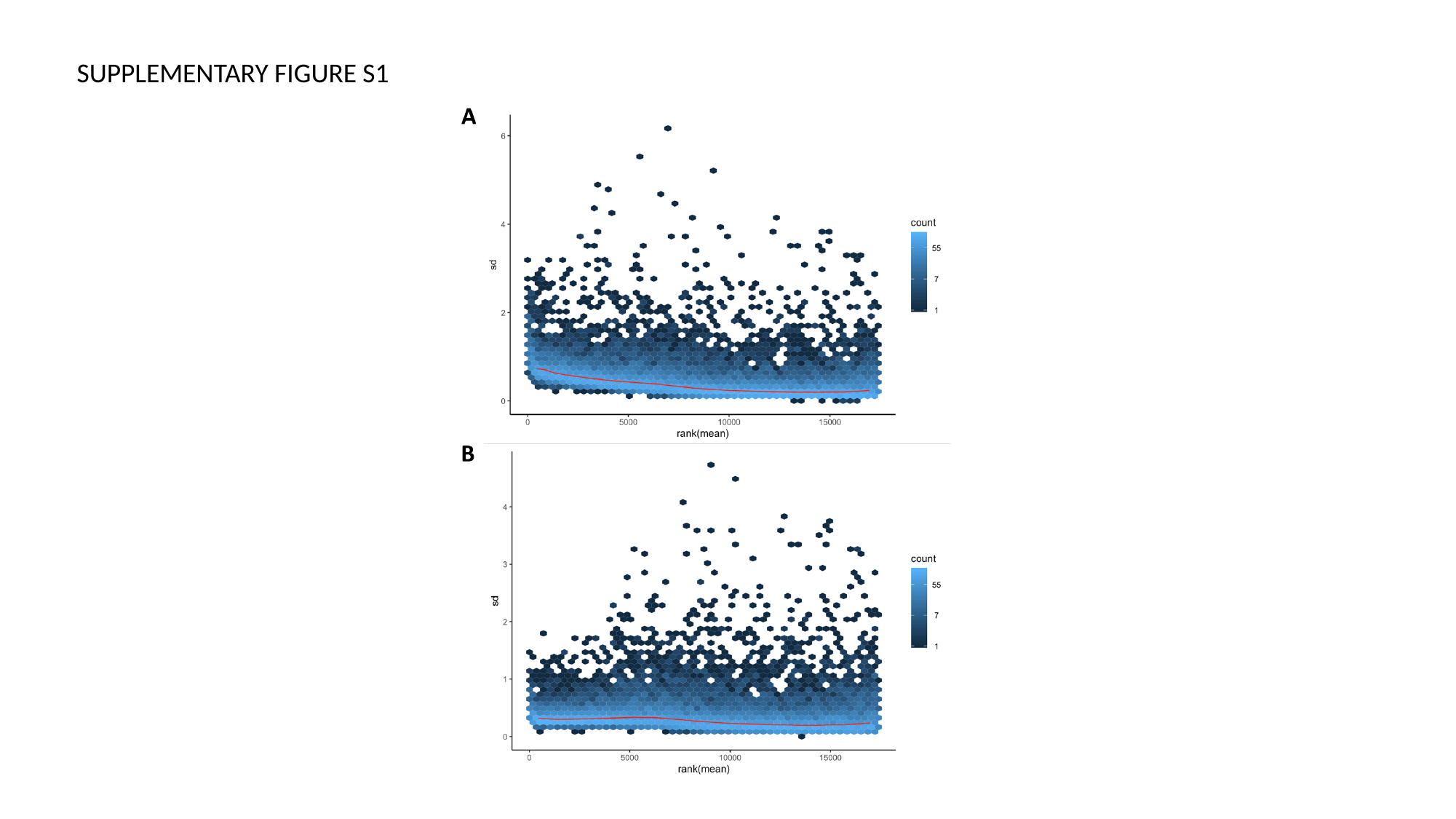

SUPPLEMENTARY FIGURE S1
